# Valproic Acid Treatment Enhances Chromosome Flexibility and Electron Transport in MCF7 Breast Cancer Cells

**DOI:** 10.1101/2024.04.08.588551

**Authors:** Tanya Agrawal, Debashish Paul, Amita Mishra, Arunkumar Ganesan, Suchetan Pal, Tatini Rakshit

## Abstract

The structural integrity of the chromosomes is essential to every functional process within the eukaryotic nuclei. Chromosomes are DNA-histone complexes essential for the inheritance of genetic information to the offspring and any defect in it is linked to mitotic errors, cancer growth, and cellular aging. Changes in the mechanical properties of a chromosome could lead to its compromised function and stability, leading to chromosome breaks. Here, we studied the changes in chromosome physical properties using metaphase chromosomes isolated from human breast cancer cells (MCF7) exposed to Valproic Acid (VPA), a known epigenetic modifier drug involved in histone hyperacetylation and DNA demethylation. Due to chromosomal structural intricacy, preparative and technical limitations of analytical tools, we employed a label-free atomic force microscopy approach for simultaneously visualizing and mapping single chromosome elasticity. Additionally, we performed electron transport characteristics of metaphase chromosomes to elucidate the effect of VPA. Our multi-parametric strategy of probing physical properties of chromosomes offers a new scope in terms of analytical tools for studying chromosomal structural changes/aberrations and associated structure-function relationships pertinent to cancer.

## Introduction

Breast cancer (BC) is the most common cancer diagnosed among women, with a 30% increase in the incidence rate over the last twenty years^1,2^. BC progression relies on genetic and epigenetic alterations; later, being reversible provides opportunities for therapeutic applications. Nucleic acid and peptide sequencing-based mass spectrometry is usually employed for mapping post-translational epigenetic modifications. Despite numerous efforts, the pathogenesis of BC is still unclear due to its biological heterogeneity, and effective therapeutic developments are challenging^3^. It is known that DNA methylation and histone post-translational modifications regulate gene expression without altering DNA sequence, which directly contributes to the tumorigenesis and progression of BC^4,5^. Hypermethylation of CpG islands of the gene promoter is catalyzed by a group of DNA methyltransferase enzymes (DNMTs, including DNMT1, DNMT2, and DNMT3), and is linked with transcriptional silencing of the concerned genes^6^. Modification of histones is controlled by a balance between histone acetyltransferases (HATs, which add an acetyl group to N-terminal lysine residues in histones) and histone deacetylases (HDACs) activities^7-9^. An imbalance between the two expressions leads to dynamic transitions in chromatin structure and is correlated to numerous cancers^9^. Several inhibitors of DNA methyltransferases and histone deacetylases are approved by the US Food and Drug Administration (FDA) as anti-cancer drugs^10^. Valproic Acid/sodium valproate (VPA) (2-propyl pentanoic acid, low–molecular weight branched-chain fatty acid), is an FDA-approved drug for epilepsy, traditionally used as an anticonvulsant^11,12^ (to prevent or treat seizures). This drug is also extensively reported to act on epigenetic dysregulations by inducing acetylation in histones and affecting the DNA methylation status in different cancers including BC^13–17^ **(Scheme 1).**

**Scheme 1:**
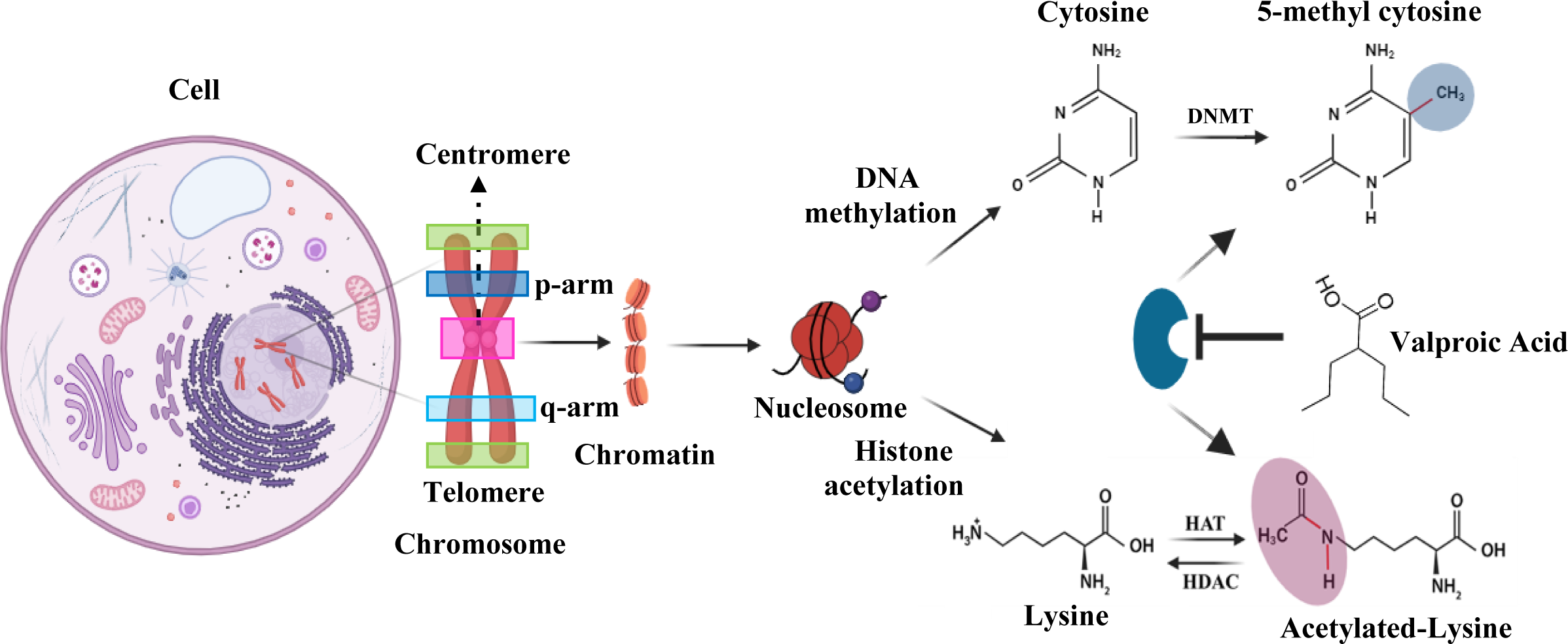
Schematic representation of the epigenetic modifications of DNA-methylation and histone acetylation in nucleosomes. Valproic acid, a well-known HDAC and DNMT inhibitor^37^. DNA methyltransferase: DNMT, Histone acetyltransferase: HAT and, histone deacetylase: HDAC.

It is known that DNA methylation and histone acetylation modulate chromatin structure. DNA methylation impacts chromatin compaction via linker DNA entry/exit angle alteration^18^. Similarly, acetylation modulates the interaction potential of the N-terminal tail domains of the core histones^19^. They both influence the folding and functional state of the chromatin fiber, which in turn modulates nucleosome accessibility to transcription machinery^18,19^. Therefore, VPA induces chromatin structural rearrangements through epigenetic reprogramming involving demethylation and acetylation of specific genes/histones^20^. Previous biochemical studies revealed that VPA prompts chromatin decondensation, repair of damaged DNA, cell differentiation, apoptosis (by upregulation of Bak, downregulation of Bcl-2 expression), and cell cycle arrest (G1 or sub-G1 phases) in BC^12,16,20–22^. Presently this compound is of great interest to the field and is undergoing many clinical trials for its anti-proliferative properties in a variety of cancer types including BC^12^. We chose MCF7 (metastatic breast adenocarcinoma, estrogen receptor (ER)-positive) for chromosome isolation, as a model cell line, that has been used for many years by multiple groups^23^. Traditional southern hybridization technique revealed that MCF7 possesses the highest hypermethylation over other commonly studied BC cell lines^24^. Also, as per recent data, VPA has significant inhibitory effects on tumor progression in ER-positive BC cell types than others^16^. Interestingly, multiple studies have revealed a higher prevalence of Chromosomal Instability (CIN) in metastatic breast cancer compared to primary breast cancer^25^. This suggests a strong association between CIN and the process of metastasis. Therefore, it is important to understand the mechanical properties of chromosome that is essential for its structural and functional integrity.

In this work, we elucidated the physical outcome of VPA treatment on MCF7 chromosomes employing atomic force microscopy (AFM), a label free single molecular detection (SMD) technique capable to provide simultaneously nanoscale resolved topography and elasticity maps (through AFM nanoindentation) at a precise location of a chromosome. The ability to study chromosomes *in situ* under physiological conditions in their pristine form is its major asset. With a conductive probe (CPAFM), AFM is uniquely capable of measuring electron flux through chromosomes with varying physical forces applied. This feature is useful in studying chromosomal structural rearrangements since cytosine methylation has a profound impact on DNA-mediated charge transport^26^. To date only a handful of reports are available on AFM-based chromosomal biophysical dissections. Lipiec et al. applied AFM-Raman to reveal molecular alterations in Hela chromosomes by inducing bleomycin^27^, previously they studied molecular differences between eu-and heterochromatin using AFM-IR based on differences in methylation status^28^. In another work, Lee group evaluated the surface charge and stiffness of Human B lymphocyte chromosomes by treating them with RNase and pepsin enzymes using KPFM and PF-QNM techniques^29^. In this study, for the first time, we quantified both the single chromosome elasticity and electron transport through isolated MCF7 single metaphase chromosomes using AFM to decipher the correlation between the chromatin structural alterations induced by VPA and its therapeutic function. This quantitative approach is valuable for assessing the disease progression, efficiency of the drug treatment, and associated adverse effects. The experimental setup is affordable and does not require special expertise to perform the required measurements and data analysis.

### Experimental

#### Cell Culture and metaphase chromosome isolation

Human BC cell line MCF7 was purchased from the National Centre for Cell Science (Pune, Maharashtra, India) and cultured according to the manufacturer’s protocol. Cell cultures were incubated at 37 °C in a humidified environment having 5% CO_2_. Metaphase chromosomes were isolated from MCF7 cells according to the protocol detailed elsewhere^30^. Briefly, 0.1 μg/mL colcemid (Sigma-Aldrich, USA) was added at 60−80% cell confluency and incubated overnight at 37 °C in a humidified atmosphere with 5% CO_2_. Then, a controlled amount of trypsin and complete media were added, centrifuged at 1000 rpm for 5 min. After that excess supernatant was discarded. Next, prewarmed KCl (Sigma-Aldrich, USA) was added, mixed well and the mixture was incubated for 20 min in a 37 °C water bath. Subsequently, 5 drops of freshly prepared fixative (methyl alcohol and acetic acid (Sigma-Aldrich, USA) were mixed 3:1, by volume/volume) was added to stop the reaction. Then the content was centrifuged, and the pellet was extracted. After washing the pellet, the chromosomal sample’s concentration was checked by Nanodrop (Thermo Fischer). The chromosomes were treated with RNase (Sigma-Aldrich, R6513) and protease (Sigma-Aldrich, P6887) to remove nucleic acid and protein contaminants^31^. The sample was kept in the fixative solution at 4 °C for future use (maximum 3 days). 1µM VPA (VPA) (Sigma-Aldrich) dissolved in MilliQ water was added to chromosome solution for 1 h before performing experiments. Chromosomes were routinely checked under an optical microscope and a fluorescence microscope (stained with acridine orange. Ti2E A1R MP, Nikon) before and after the VPA treatment. Images were processed and quantified using Image J software.

#### Fourier-transform infrared (FTIR) measurement

FT-IR spectra of chromosome samples were taken using the KBr window in Nicolet iS20 FTIR spectrometer (Thermo Fisher Scientific, Massachusetts, United States). The infrared signal was measured by a deuterated triglycine sulfate (DTGS) detector. Each spectrum was an average of 64 scans in the range between 400 and 4000 cm^−1^, with 4 cm^−1^ resolutions.

#### Circular Dichroism (CD) Measurement

Circular dichroism spectra of chromosome samples were taken using a Jasco J-1500 CD spectrophotometer at 4 °C. The deconvolution of the CD signals into the relevant secondary structure was carried out by CDNN software, which was evaluated using the BestSel CD deconvolution server.

#### PCA Analysis

Here, spectral data for each chromosome sample were collected for multivariate analysis. Baseline correction and vector normalization were achieved for each spectrum separately using Origin Lab (Northampton, MA). The first derivative was taken to minimize the baseline error and then smoothed by using the Savitzky−Golay method, and finally, PCA analysis was done by using Origin Software.

#### UV-Vis Measurement

The UV−visible absorption spectra were collected using a Shimadzu UV-2600i/2700i spectrophotometer (Kyoto, Japan) using a quartz cuvette having a 10 mm light path length (Singapore). The optical direct band gap was estimated from the (αhν)^2^ vs hν curve (Tauc plot), where α, h, and ν are the absorption coefficient, Planck constant, and light frequency, respectively.

#### Electrochemical measurements

A three-electrode cell comprising a glassy carbon working electrode, Ag/AgCl in 1 M KCl/1 M KNO_3_ reference electrode, and a platinum wire counter electrode, was used (Bio-Logic SP-300 Potentiostat at 25 °C, PBS used as electrolyte solution). Chromosome samples (untreated and VPA-treated) in PBS buffer (pH ∼ 7.4) were deposited on the electrode, the signals were recorded by scanning the potential between −1.5 and +1.5 V using cyclic voltammetry with scan rates of 25, 50, 100, and 200 mV s ^−1^. Amplitude; 10 mV, frequency range; 7MHz to 100mHz was applied for the Electrochemical Impedance Spectroscopy (EIS).

#### AFM imaging and analysis

For AFM sample preparation, freshly prepared chromosomes solution (untreated and VPA-treated MCF7 chromosomes) was placed on a cleaned glass slide and incubated for 20 min, washed briefly with PBS. Fluid imaging (in intermittent contact mode) and force spectroscopy were performed on an Asylum MFP-3D instrument (Asylum Research, Santa Barbara, USA) with a triangular silicon nitride probe (spring constant 0.3 N/m) under PBS buffer (pH 7.4). All AFM images were analysed using the Igor Pro software.

#### AFM nanoindentation

The chromosome spread was imaged in PBS buffer in intermittent contact mode. Each chromosome and sub-chromosomal region were zoomed in and reimaged before starting force spectroscopy. Cantilever spring constant (0.1-0.3 N/m) and deflection sensitivity values were calibrated before each experiment. We carefully checked the height values of all different parts of a single chromosome from fluid AFM images. The height values were always greater than 85 nm. To generate force maps, an area of 500 x 500 nm was selected on a single chromosome using a maximum applied force of 2 nN. Maximum 10-15 nm of indentation depths were made. The Young’s modulus values were determined from the force maps using the Hertz model with spherical geometry having the equation δ=[3(1-ν^2^)/(4ER^1/2^)]^2/3^F^2/3^. Here, δ, F, E, and R are the indentation, force, Young’s modulus of the sample, and radius of the tip respectively, and ν is the Poisson’s ratio of the sample which is assumed to be 0.3^32^ using Igor Pro software.

#### CP-AFM analysis

An Indium tin oxide (ITO)-coated glass substrate was thoroughly cleaned^33^. Freshly prepared chromosome solutions were deposited on a cleaned and dried ITO (1.0 cm^2^) substrate, washed briefly with PBS and dried with a weak stream of nitrogen gas. Each sample was imaged by intermittent-contact AFM (MFP-3D Asylum Research, Santa Barbara, USA using AC240TS-R3 (Oxford Instruments, USA)) to check the position of chromosome spread. For I-V measurements Pt–Ir-coated silicon cantilever (HQ-DPER-XSC11, MikroMasch, with a quoted radius of 20nm) was used. Potential sweeps were made from -3V to +3V and the resulting current responses were recorded (MFP-3D -ORCA mode). Each I-V spectrum acquired on the single chromosome was the average of 4 sweeps. During each measurement, the force set point (10-100 nN) was calibrated and adjusted by applying a definite set point^34^. All CSAFS measurements were performed in ambient conditions where temperature and humidity were maintained at 24 ± 1 ^0^C and 35–45%, respectively. 100 I–V curves for both untreated and VPA-treated chromosomes at 10, 40, and 80nN force loads were taken after careful selection and averaged. The *I*–*V* curves for both systems at different force loads were fitted with the Fowler–Nordheim equation using Origin 8.5 software^34^.

#### ICP-MS analysis

Freshly prepared chromosome solution was put in freshly prepared aqua regia (1:3 conc. nitric acid and hydrochloric acid) and heated for 12 h at 60 °C for complete digestion, analyzed with an Perkin Elmer inductively coupled plasma mass spectrometer (ICP-MS)^35^.

## Results and discussion

### Spectroscopic investigations on VPA treated MCF-7 chromosomes

The research involving various BC cell lines revealed that cellular response to VPA treatment depends on the type of cell, drug dosage and time^12,17^. In MCF7 cell, VPA was reported to induce apoptosis at the lowest dose of 0.5mM after 8 days of treatment^36^. In a similar study, Masumeh et al. reported a VPA -IC50 value of 3µM^21^. Earlier reports also found morphological changes in MCF7 cells after exposure to VPA for more than 48 hrs^16^. In this report we performed spectroscopy experiments to confirm the VPA treatment to MCF7 cells prior to start AFM measurements. We varied the concentration of VPA and incubation time to find out the lowest concentration (1 µM) and minimum incubation time (1hr) to get reproducible and notable changes in spectroscopic signatures.

We first attempted IR spectroscopy for untreated and VPA treated chromosome samples. The major differences were observed in the spectral range of 800-1800cm^-1^ and 2850-3000 cm^-1^ for both untreated and VPA-treated chromosomes (**Figure 1, A-D**). 2850-3050 cm^-1^ was assigned to ν_sym_ and ν_asym_ stretching of -CH_2_, -CH_3,_ and C-H vibrations in 5-methylcytosine methyl groups. The increase in absorbance intensity was observed for untreated MCF7 chromosomes compared to treated sample^30^. Corresponding area under the curve (AUC) values were also calculated (which signifies the absorbed energy)^37^. The band peak area for untreated chromosomes was ∼1.23 times higher than VPA-treated chromosomes (**Table S2**)^30^. The band centers were found at 2925 and 2923cm^-1^ in untreated and VPA treated ones respectively. The peaks∼1070cm^-1^ was assigned for C-O stretching and −PO_2_ symmetric stretching (**Figure 1B**), ∼1240 cm^−1^ for nucleobases and −PO_2_ antisymmetric stretching of the DNA backbone (related to conformational changes) were noticed (**Figure 1E**). The absorption intensity of untreated sample was lower than VPA treated chromosomes due to a higher abundance of cytosine methylation^38^. Untreated and VPA-treated chromosomes consisted of peaks at 1388cm^-1^, which corresponded to the -CH_3_ deformation of 5-methylcytosine and the vibration of cytidine. The peak at 1755cm^−1^ corresponded to the C=O stretching of guanine, which was observed in both cases. The spectral range between 1610−1695 cm^−1^ was of Amide I (predominantly C=O stretching) of histone and C=O stretching, out-of-plane −NH_2_ bending, dT, dG, dC, C/G/T, C=C stretching, and in-plane −NH_2_ bending of DNA, changes in the intensity of peaks in this region represented the histone modifications (**Table S1**). The spectral range of 1200-1300cm^-1^ represented amide III region, where higher intensities observed for VPA-treated chromosomes signifying loosely packaged chromatin^27^ (**Figure 1E**). The peak ∼1171cm^-1^ was for asymmetric CO stretching of bridge oxygen (ν_co_). We observed an intensity difference between the two cases suggesting different levels of acetylation^39^. (**Figure 1E**). Next, we performed principal component analysis (PCA) (400-4000cm^-1^) which clearly displayed the distinctions between untreated and VPA-treated chromosomes (**Figure 1F**). PC1 and PC2 components provided the dominant account for the maximum variance in the data set. The overall analysis from FT-IR spectra and PCA analysis deciphered that VPA treatment to chromosomes reduced DNA methylation and elevated histone acetylation levels. To analyze the effect of CpG methylation and histone acetylation, we performed circular dichroism (CD) measurements (**Figure 1G**). Through % secondary structure evaluation, we observed helix, antiparallel beta-sheet and turns (190-220 nm) were preferentially found in VPA-treated chromosomes than untreated ones (**Table S3**), this is linked to increased acetylation of histones with VPA^40,41^. The spectral range of 260-300nm represented the CpG methylation^42^. We also performed CD-PCA (190-300 nm) (**Figure 1H**) that clearly distinguished the untreated and VPA treated chromosomes.

**Figure 1:**
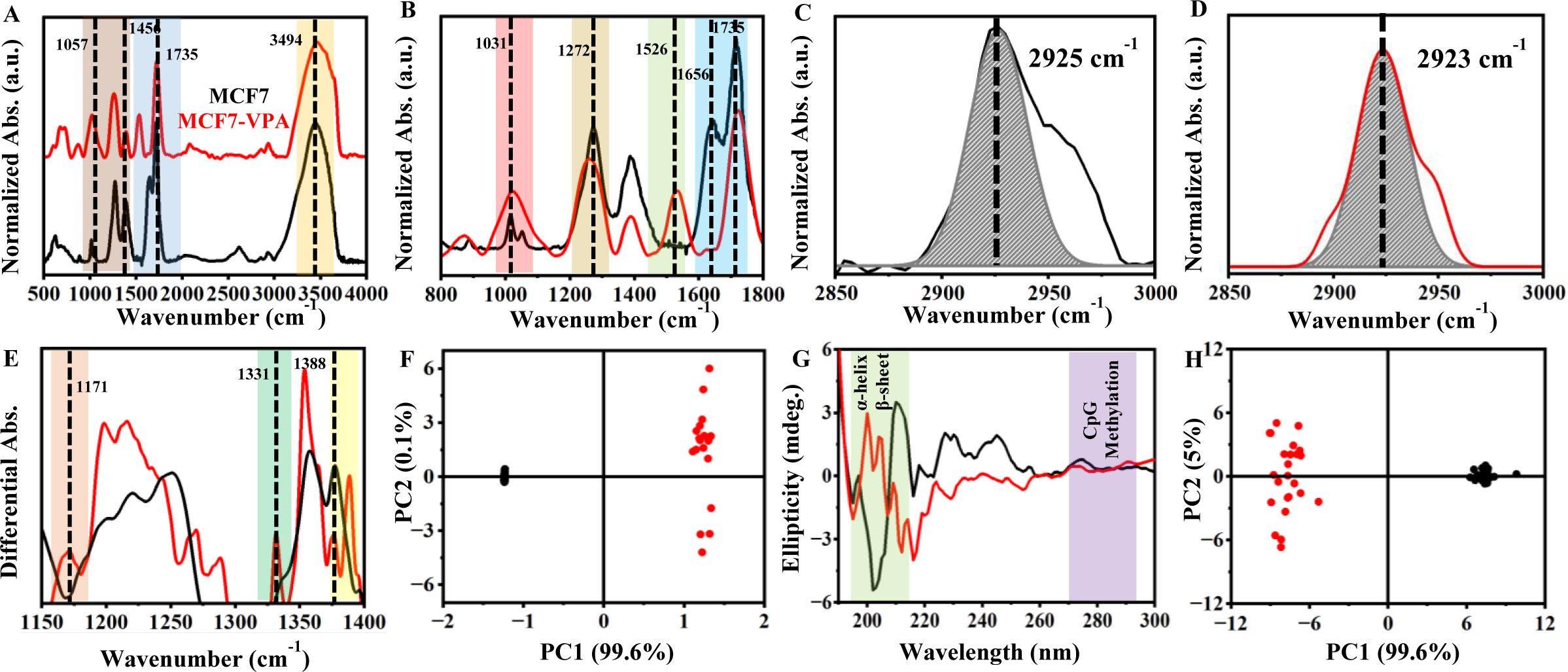
(A) FT-IR average spectral profile of MCF7 (black) and MCF7-VPA treated (red) chromosomes. (B) Mid-infrared spectral overlay of untreated and VPA-treated chromosomes, which represented different spectral signatures due to different methylation status in the wavenumber range 800−1800cm^−1^. Area under the curve (AUC) after Gaussian peak fitting for (C) MCF7 (D) MCF7-VPA treated chromosomes, which represented the DNA−CH_3_ band. (E) Savitzky−Golay’s second-derivative spectra in the wavenumber range of 1150−1400 cm^−1^ showing the degree of acetylation for the two samples. (F) PCA score plot of both chromosomes. The scores are based on the first two principal components: the region used for PCA was 400−4000 cm^−1^. (G) Circular dichroism spectra (H) PCA score plot of both samples

### AFM Nanoindentation based mechanical characteristics of chromosomes

We visualized MCF7 chromosomes with high resolution AFM in intermittent contact mode under physiological buffer conditions (**Figure 2**). Frequently, ultrastructural components of centromere and telomere were also discerned (**Figure 2C)**. We routinely checked the chromosome preparation with light/fluorescence microscopy before AFM imaging (**Figure S1**). We visualised chromosomes with VPA treatment *in situ* and compared the height and roughness values of both untreated and treated chromosomes **(Figure 3, A-C)**. Both the measured values were significantly changed before and after VPA treatment (**Figure 3, D-E**), height values were decreased in 4 chromosomal regions (145 nm (untreated) vs. 134 nm (VPA -treated)) and roughness values were increased (18 nm (untreated) vs. 31 nm (VPA -treated)), these observations are linked to VPA-induced structural alterations in treated chromosomes. (**Table S4 and S5**).

**Figure 2:**
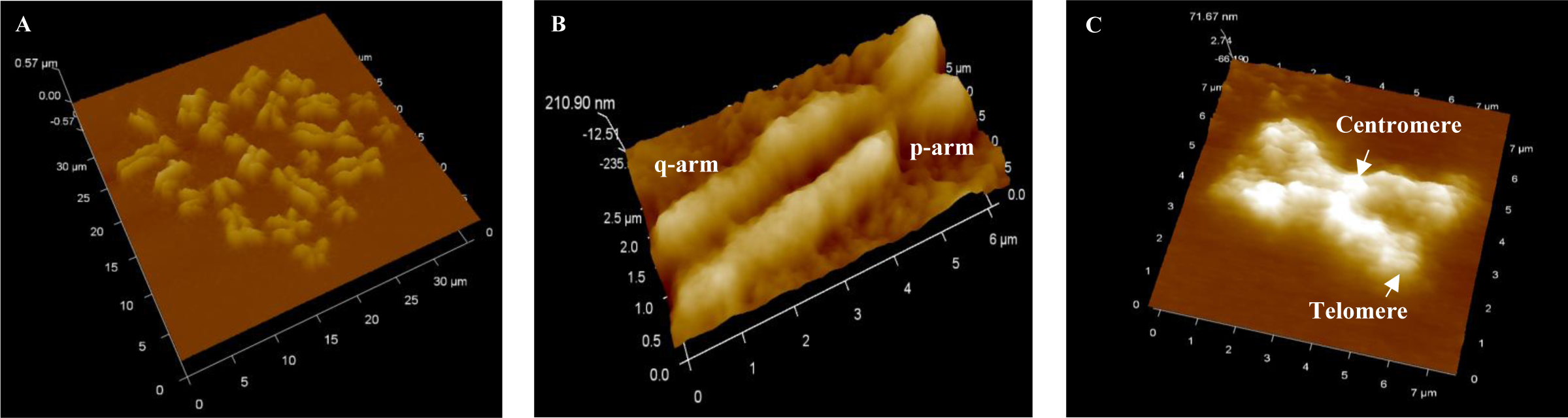
AFM topographic analysis of the MCF7 chromosomes in buffer (A) 3D AFM topographic image of a chromosome spread (B) single chromosome, (C) chromosome ultrastructure

**Figure 3:**
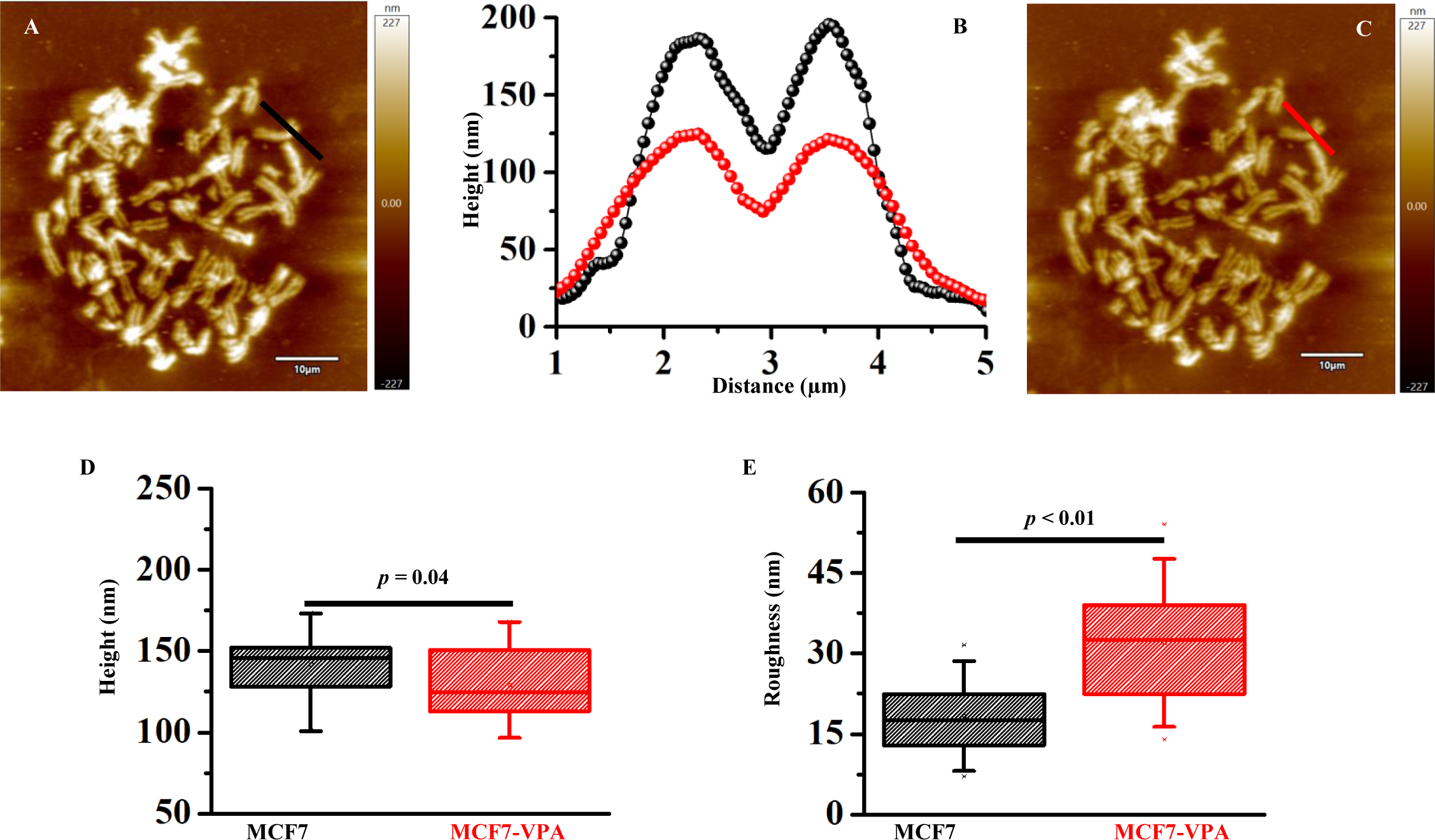
AFM analysis of VPA treatment process of MCF7 chromosome (A) untreated and (C) VPA-treated chromosome spreads. Data were obtained with the same chromosome set by treating with VPA *in situ,* (B) Cross-sectional profiles corresponding to the lines in the AFM images in (A and C). Box plots of (D) height and (E) surface roughness (Ra) values of untreated and treated chromosomes ( *p* values from unpaired t-test are shown).

Next, AFM nanoindentation force spectroscopy was employed under physiological buffer conditions to measure the elasticity of a single chromosome. AFM nanoindentation mode provided a force-distance curve expressing the force exerted (max. 2nN) on a single chromosome by the AFM tip as the tip indented (max. 20 nm) the chromosome. Young’s modulus was obtained by fitting the approach curve with the Hertz model (spherical indenter geometry) for both cases. (**Figure 4A**)^32,43,44^. We found that the VPA-treated chromosomes were strikingly 6 times more elastic than untreated chromosomes (12.5±2 MPa (untreated) vs. 2.0±0.6 (VPA-treated), p<0.01) (**Figure 4, F, I**). The range of Young’s modulus values of chromosomes were consistent with previous reports^29,45^. This data reveals how epigenetic reprogramming impact material properties of chromosomes. DNA methylation is the addition of a methyl group to cytosine DNA bases (**Scheme 1**), which decreases the repulsion between DNA strands, resulting in a more compact chromatin structure^46^. Earlier reports suggested that DNA demethylation can open up chromatin and prevents the strands from becoming tangled and also plays important roles in reinforcing the DNA during cell division, preventing DNA damage, and regulating gene expression and DNA replication^47^. In case of hyperacetylation of histones, it relaxes chromatin structure by adding negatively charged acetyl groups to specific lysine residues in histone (**Scheme 1**). This decreases the electrostatic affinity between histone proteins and DNA, which disrupts their interactions. The result is a more relaxed chromatin structure, which is associated with greater levels of gene transcription. Histone hyperacetylations are critical to various cellular processes, including nucleosome assembly, chromatin folding, and DNA damage repair. ^48,49^. Therefore, VPA transformed the condensed chromatin to its relaxed form (**Scheme 2**). We rigorously captured AFM force-volume maps at different sub-chromosomal regions (centromere, telomere, p, and q-arms) for mapping elasticity (**Figure 4B-C**) to investigate the effect of VPA on it. We found that VPA-treated chromosomes were uniformly elastic in 4 chromosomal regions than untreated ones. (**Table S4**, **Figure 4, D, G**). The average height, Young’s moduli and two-tailed unpaired t-test results of different sub-chromosomal regions are represented in **Table S4 and S5**.

**Figure 4:**
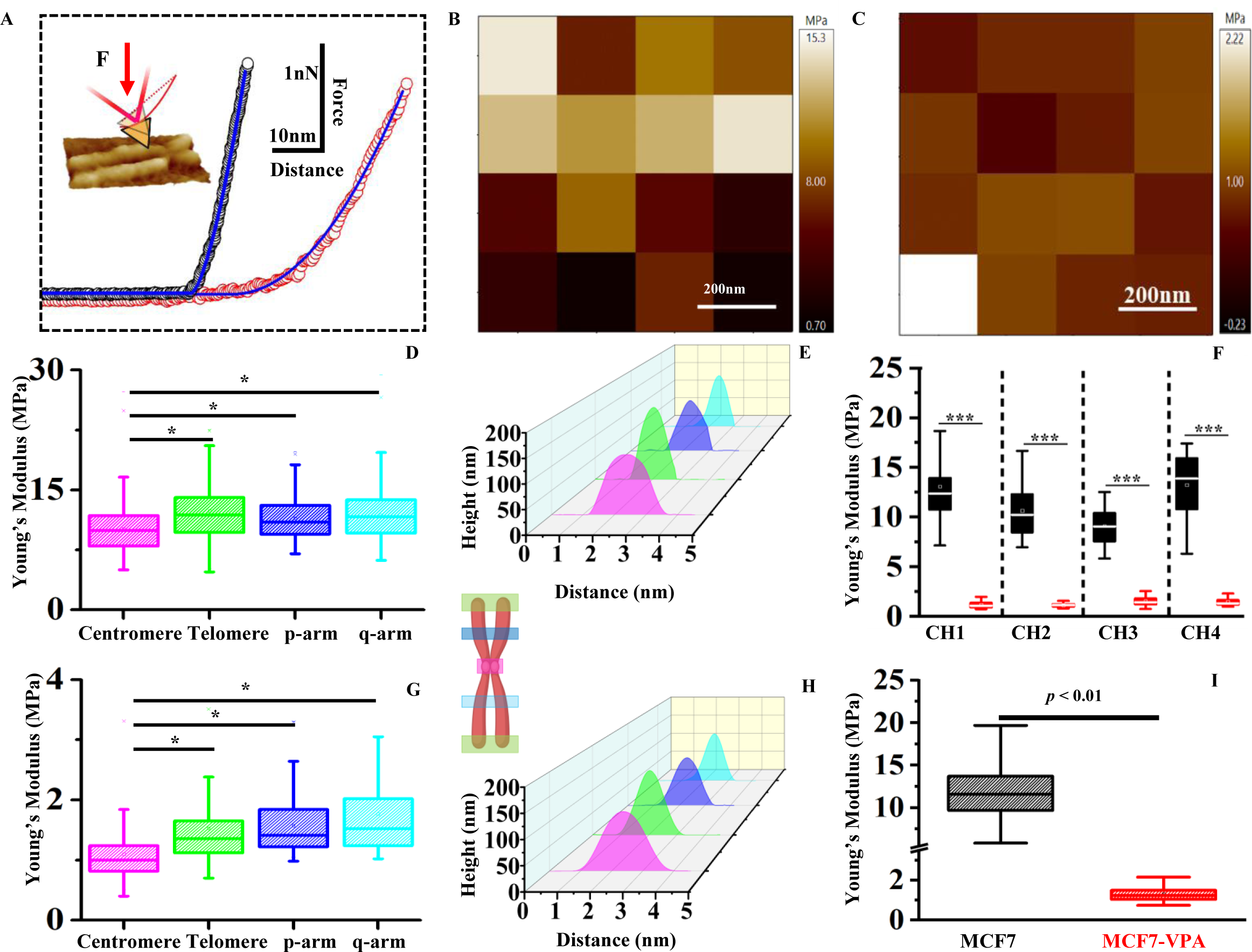
(A) A sample force–distance profile representing AFM nanoindentation on a single chromosome of both MCF7 (black) and MCF7-VPA treated (red), fitted with Hertz model with spherical indenter geometry, a 3D topographic view of chromosome on a glass slide under buffer (inset). The force maps of Young’s modulus for (B) MCF7 and (C) MCF7-VPA treated chromosomes at the centromere. The Box plots of Young’s modulus (using Hertz model) values and height distribution profile for 4 different (D-E) MCF7 and (G-H) MCF7-VPA treated chromosomes at different chromosomal regions, (F) Comparison of average Young’s modulus values of four MCF7 and MCF7-VPA treated chromosomes (considering all 4 regions), (I) Average Young’s modulus of untreated and treated chromosomes, *p*-values were calculated by one-way ANOVA, followed by the Tukey posthoc test.

**Scheme 2:**
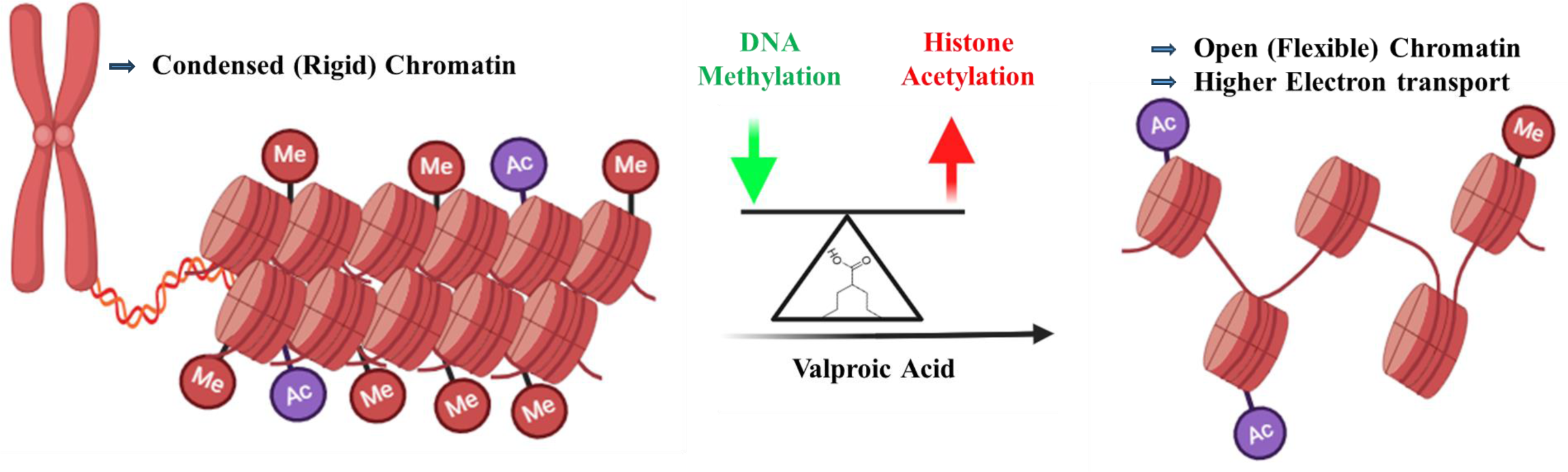
Changes in DNA methylation and histone acetylation levels after VPA transforms condensed chromatin to its open conformation.

### CPAFM study to assess electron transport through chromosomes

We performed electrical measurements at the bulk level to analyze the effect of VPA incubation with MCF7 chromosomes by Electrochemical Impedance Spectroscopy (EIS)^50^, cyclic voltammetry (CV) and UV-Vis-Tauc plot ^30^. A consistent trend was observed between untreated and VPA treated chromosomes in all the three modes of operation (**Figure S2, S3**), where the latter was found to be more conductive. These current sensing analyses provided crucial insights into the electronic capabilities of the chromosomes at the bulk level and prompted us to check the electron transport through single chromosomes.

We performed CPAFM analysis to quantify electron transport through untreated and treated single chromosomes. In CP-AFM, the conductive probe physically touches the biomolecule, the tip positioning is decoupled from the sample conductivity allowing more precise knowledge of the relative positions of the tip and sample in the vertical plane. Electron transport characteristics of biological macromolecules can be assessed with carefully maintained varying applied forces^51^ (**Figure 5A**). Freshly prepared chromosome samples were deposited on ITO substrates for CPAFM and current–voltage (I-V) curves were captured under ambient conditions. The I-V characteristics of both untreated and VPA treated chromosome samples were captured at ±3V applied bias with 10, 40, and 80nN forces. Chromosomes are supramolecular complex of DNA and proteins. They were conductive due to the presence of DNA, proteins^31,52^ and cupric ions bound to H3-H3ʹ interface of the histone tetramer^53^ (**Figure 5B**). The measured conductivity was comparable to metalloproteins^34^. We quantified cupric ions with ICP-MS measurements, 4x 10^4^ ppb copper were found to be associated with one preparation of MCF7 chromosomes where we started with approximately 10^6^ cells.

**Figure 5:**
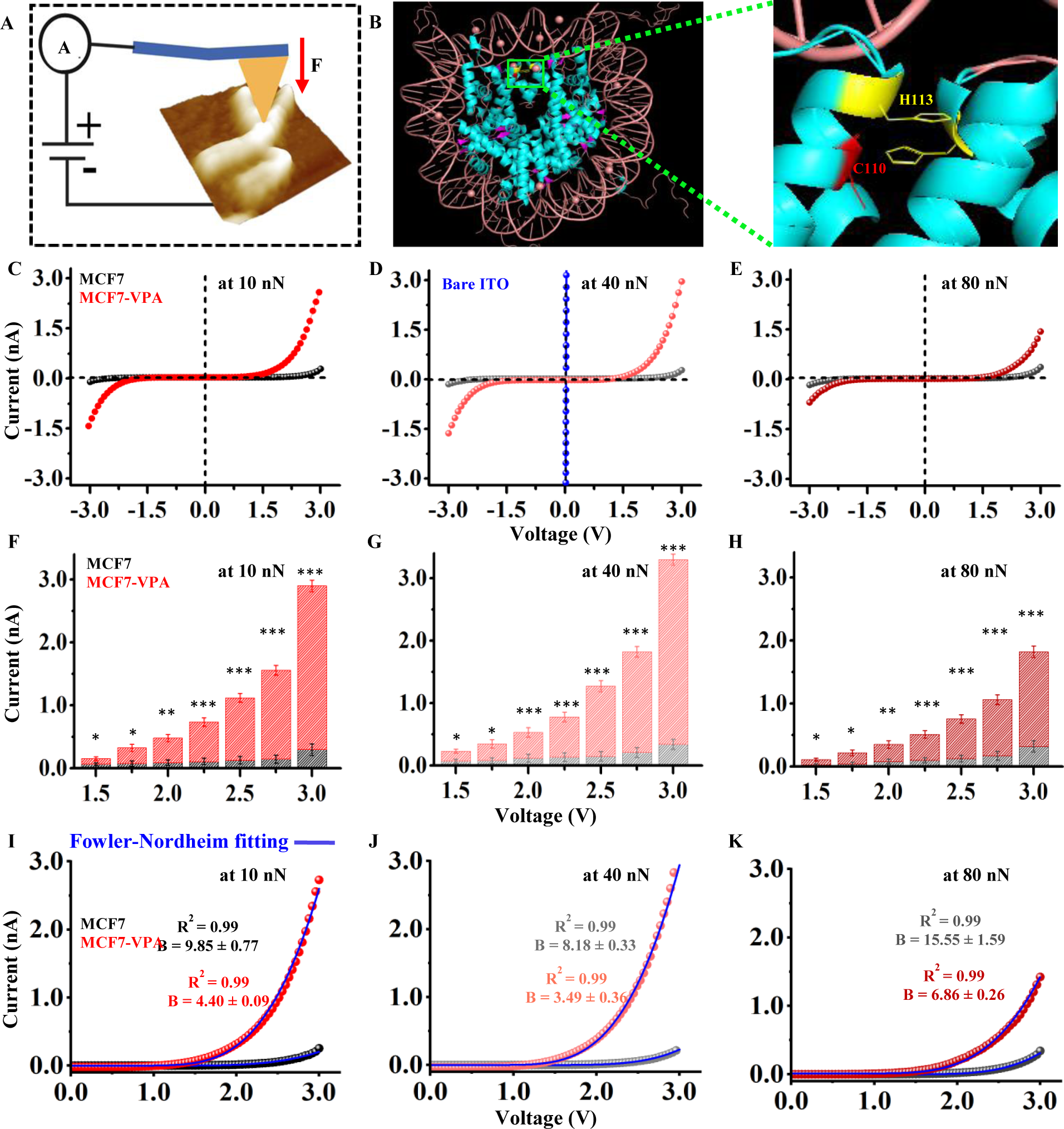
(A) Schematic illustration of junction configuration employed in the ETp measurement via single chromosome on an ITO substrate (B) Nucleosome core particle structure (Protein Data Bank (PDB) 1KX5). The box represents the H3-H3’ interface. Interface residues H3H113 and H3C110 are shown^51^. I-V responses for bare ITO (blue), untreated, and VPA-treated chromosomes over the bias range of ± 3.0V at (C) 10 (D) 40 (E) 80 nN applied forces (tip-positive). (F-H) The histograms showing the current values against applied voltage for both chromosomes at 10, 40 and 80 nN forces. The overlay of I–V curves at (I) 10 (J) 40 (K) 80 for MCF7 and MCF7-VPA treated chromosomes fitted with Fowler–Nordheim (F–N) model (blue line) of the form I(V) = AV^2^ exp (-B/V) for obtaining the respective ‘‘B’’ values^33^.

At all the applied forces, VPA treated chromosomes were more conductive than untreated samples, and the current values were significantly higher than untreated chromosomes shown in histograms (**Figure 5, C-H**). With increasing applied forces from 10-40nN, the current values increased for both samples. However, we observed a decrease for both at 80nN, the underlying reason was not apparent. We found that our measured I-V curves could be well fit with Fowler–Nordheim (F-N) tunneling model of the form, I(V) = AV^2^ exp(-B/V)^34^. This model was only applicable when the barrier height between the underlying ITO substrate and chromosomes was low. According to the F-N equation, higher the conductivity, lower the B value (details of B value calculation^54^). To calculate the corresponding “B” values, the positive side of the curves was used, as the CP-AFM tip was positive relative to the substrate (**Figure 5I-K**). The B values for untreated chromosomes was found to be more than 2 times higher than VPA-treated chromosomes under all forces. (**Table S6**). Therefore, electron transport was facilitated more in VPA-treated chromosomes than untreated ones.

It is well reported that the presence of methyl group impacts the DNA duplex stability and charge transport pathways^55^. Tao et al. observed a significant change in conductance upon methyl modification on cytosine. The destabilization due to cytosine methylation led to increased HOMO-LUMO gap, which ultimately affected the electron transport characteristics of methylated DNA^30,52,56^. Earlier report also suggested that electron transfer is highly facilitated from histone protein to DNA (thymine bases) in irradiated damaged chromatin^57^. In our study, epigenetic alterations resulted in an open relaxed chromatin (**Scheme 2**) that highly impacted DNA-histone interactions, provided better accessibility to cupric ions and significantly enhanced the conductivity of the VPA treated chromosomes.

## Conclusion

VPA induced demethylation of DNA and hyperacetylation of histones, both factors together transformed the undulant condensed MCF7 chromatin to its relaxed form. The chromosomal structural alterations resulted in increased resilience and electron transport capabilities of VPA treated MCF7 chromosomes. Chromosomal instability has a substantial impact on the growth and advancement of cancer. A multitude of variables can induce abnormalities in chromosomes, including epigenetic dysregulation. Chromosomes have inherent mechanical qualities that are crucial for their optimal functioning. Changes in these mechanical properties can significantly impact the stability and functioning of chromosomes. Therefore, the mechanical data of cancer chromosomes has a lot of implications on how the cancer therapeutic drugs works at the chromatin level, mainly because it could help assess chromosomal instability in different sub-parts of the chromosomes in the future. Tumor treating fields (TTF) combined with chemotherapeutic drugs are increasingly used for treating aggressive brain tumours, a few clinical trials are undergoing^58^. TTF is a cancer treatment that uses alternating electric fields to disrupt the ability of some types of tumor cells to grow and spread by targeting proteins in cancer cells that are essential to cell division. The electron flux data holds insight on how chemotherapy drugs combined with tumor treating fields^59^ could be manipulated for better therapeutic outcome in aggressive cancers. Also, chromosome-based bioelectronics for medical device applications is not explored so far. We are hopeful in applying multiparametric AFM and statistical (multivariate) analysis together to reveal the intrinsic connections between chromosomal structural aberrations and functional outcome.

## Author contributions

Methodology, validation, formal analysis, data interpretation, investigation, writing – original draft, visualization: T. A., D. P., A. M., Conceptualization, data interpretation, writing and editing: T.R., S. P., A.K.G. Conceptualization, methodology, writing and editing, supervision, project administration, funding acquisition: T.R., S. P.

## Acknowledgements

T.R. thanks SERB (CRG/2019/007013) and Shiv Nadar Institution of Eminence, Delhi NCR for funding and research facilities including MFP-3D AFM, and S.P. thanks IIT Bhilai, CG, SERB (CRG/2023/003029), and DBT (BT/12/IYBA/2019/14).

## Table of Content

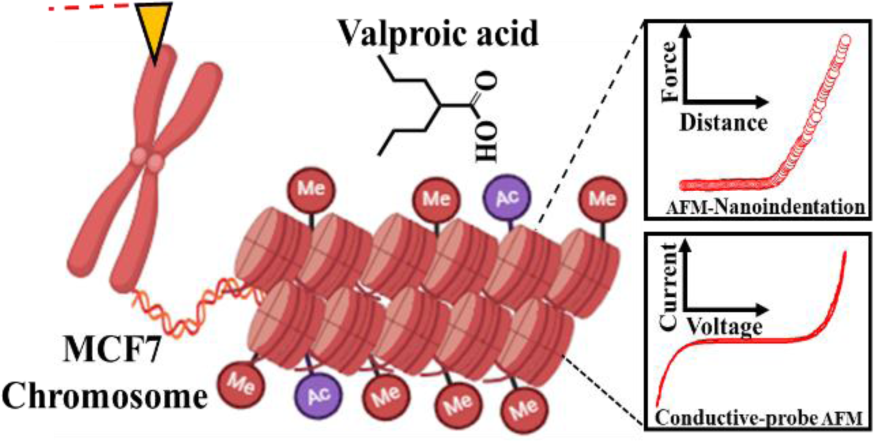

## Supporting Information

**Figure S1:**
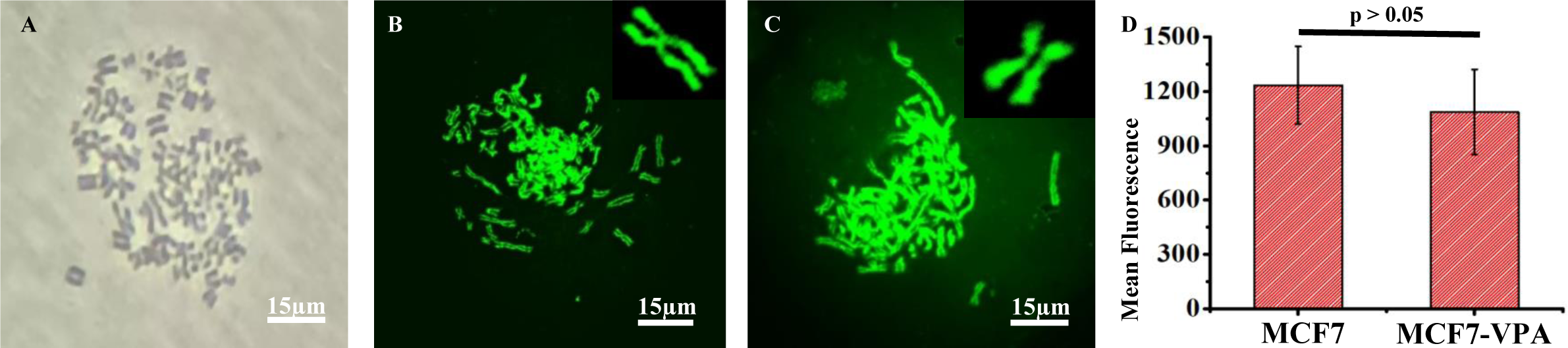
(A) Optical and Fluorescence microscopy of (B) MCF7 (C) MCF7-VPA treated chromosomes. (D) Mean fluorescence intensity plot of untreated and VPA-treated MCF7 chromosomes (Acridine orange was used for staining the chromosomes).

**Figure S2:**
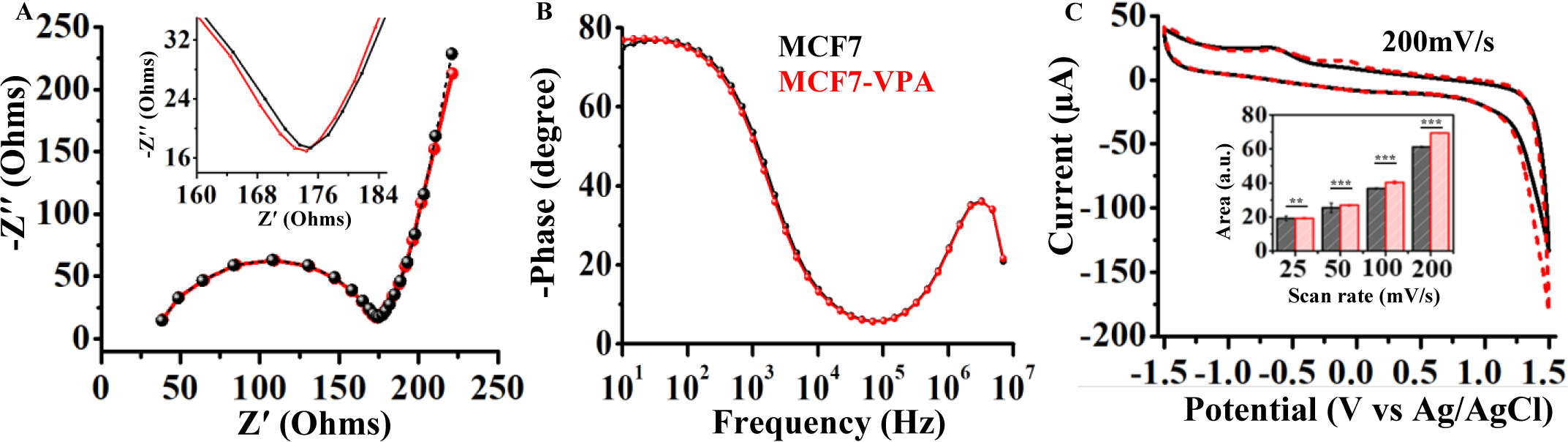
(A) Electrochemical characterization (A) Nyquist plot (B) Bode plot for both untreated and VPA-treated MCF7 chromosomes. (C) Cyclic voltammogram (CV) in PBS buffer (pH-7.4) at 200mV/s scan rate. The area under the curve (AUC) calculated from the CV voltammograms for both untreated and VPA-treated MCF7 chromosomes (inset) suggests the changes in the charge transport ability, with results expressed in mean ± standard variation (n = 3) (*** means p < 0.05 obtained from unpaired students t-tests).

**Figure S3:**
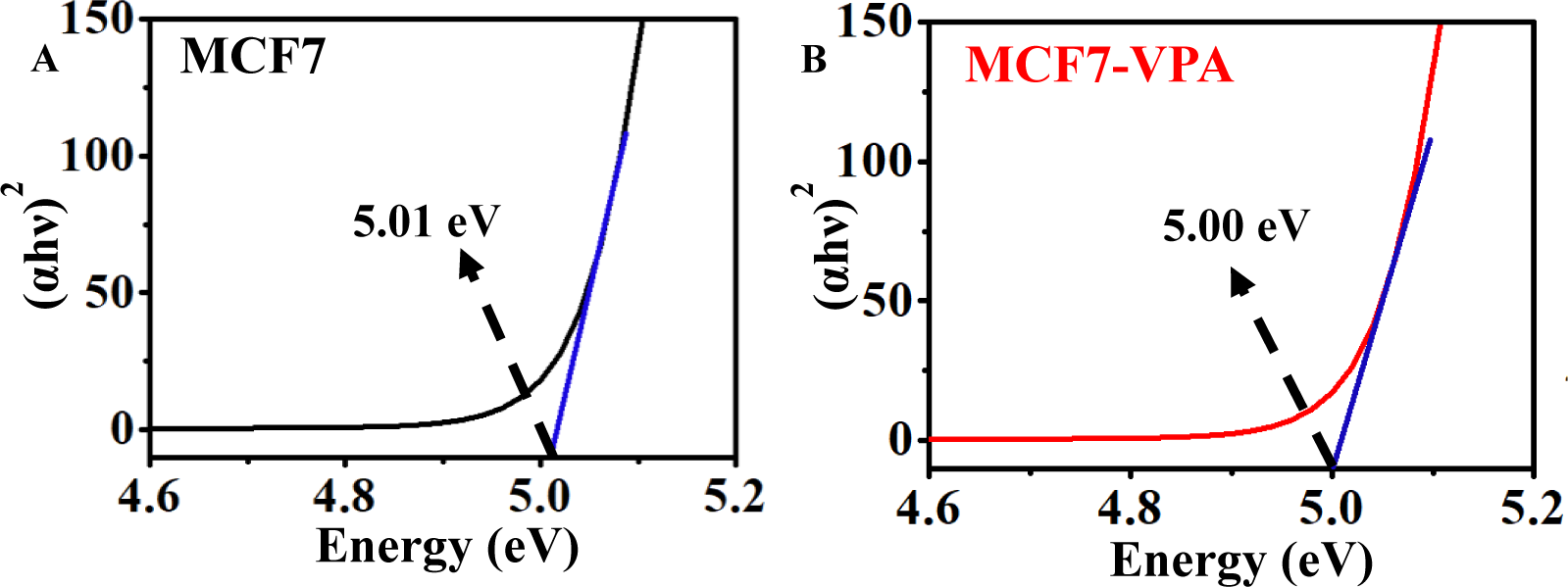
Tauc plots of (A) MCF7 and (B) VPA treated MCF7 chromosome samples obtained from the UV–vis spectra.

**Table S1:**
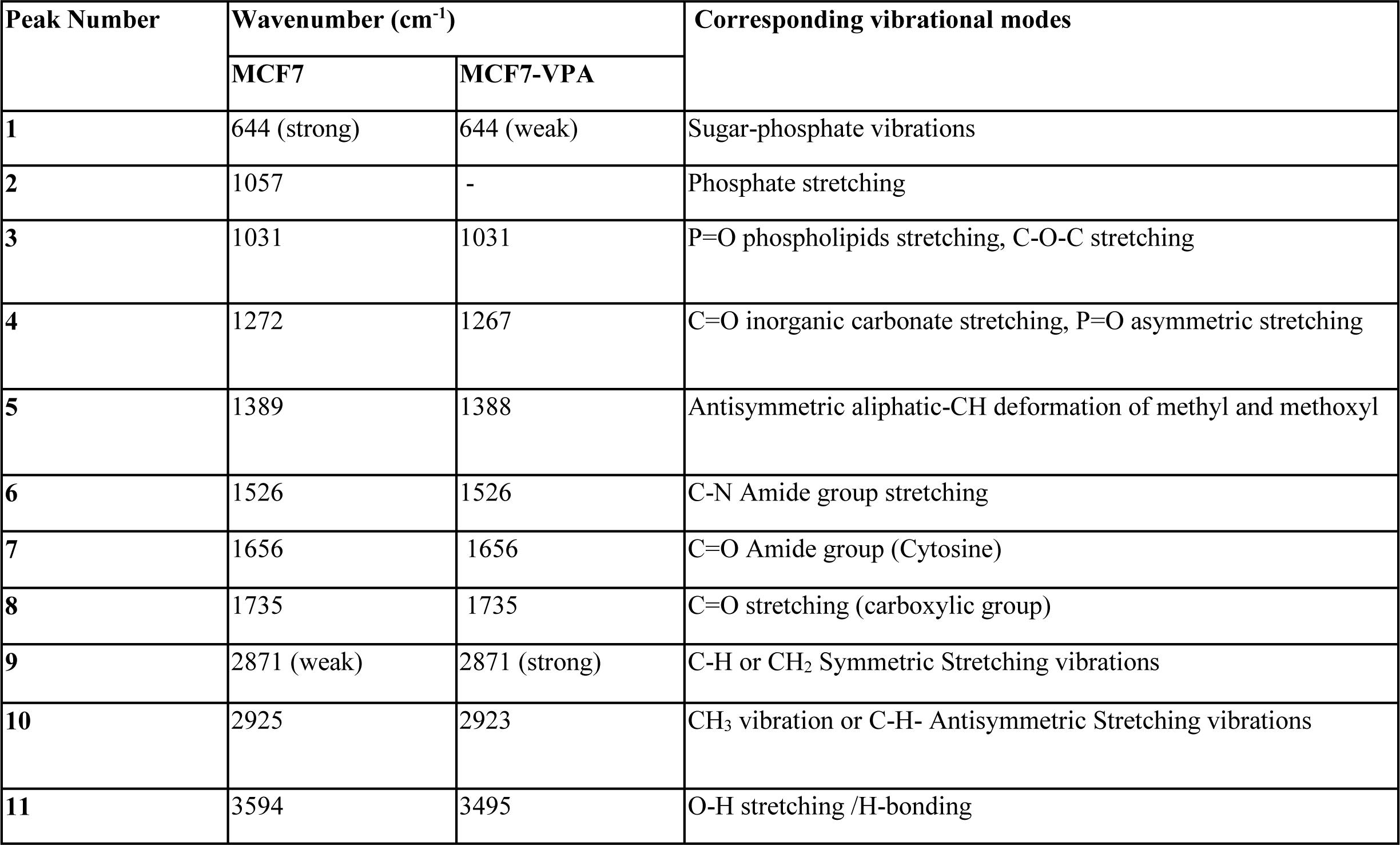
FT-IR Peak Assignments.

| Peak Number | Wavenumber (cm<sup>-1</sup>) |  | Corresponding vibrational modes |
| --- | --- | --- | --- |
|  | MCF7 | MCF7-VPA |  |
| 1 | 644 (strong) | 644 (weak) | Sugar-phosphate vibrations |
| 2 | 1057 | - | Phosphate stretching |
| 3 | 1031 | 1031 | P=O phospholipids stretching, C-O-C stretching |
| 4 | 1272 | 1267 | C=O inorganic carbonate stretching, P=O asymmetric stretching |
| 5 | 1389 | 1388 | Antisymmetric aliphatic-CH deformation of methyl and methoxyl |
| 6 | 1526 | 1526 | C-N Amide group stretching |
| 7 | 1656 | 1656 | C=O Amide group (Cytosine) |
| 8 | 1735 | 1735 | C=O stretching (carboxylic group) |
| 9 | 2871 (weak) | 2871 (strong) | C-H or CH<sub>2</sub> Symmetric Stretching vibrations |
| 10 | 2925 | 2923 | CH<sub>3</sub> vibration or C-H- Antisymmetric Stretching vibrations |
| 11 | 3594 | 3495 | O-H stretching /H-bonding |

**Table S2:** Statistical information for DNA-CH_3_ band peak and area under the curve values for both MCF7 andMCF7-VPA treated chromosomes obtained from FT-IR spectral profiles.

| Chromosomes | Peak starts<br>(cm<sup>-1</sup>) | Peak ends<br>(cm<sup>-1</sup>) | Area units | Ratio<br>control/treated | Peak center<br>(cm<sup>-1</sup>) |
| --- | --- | --- | --- | --- | --- |
| MCF7 | 2875 | 2975 | 0.37899 | 1.23 | 2925 |
| MCF7-VPA<br>treated | 2879 | 2967 | 0.31003 |  | 2923 |

**Table S3:**
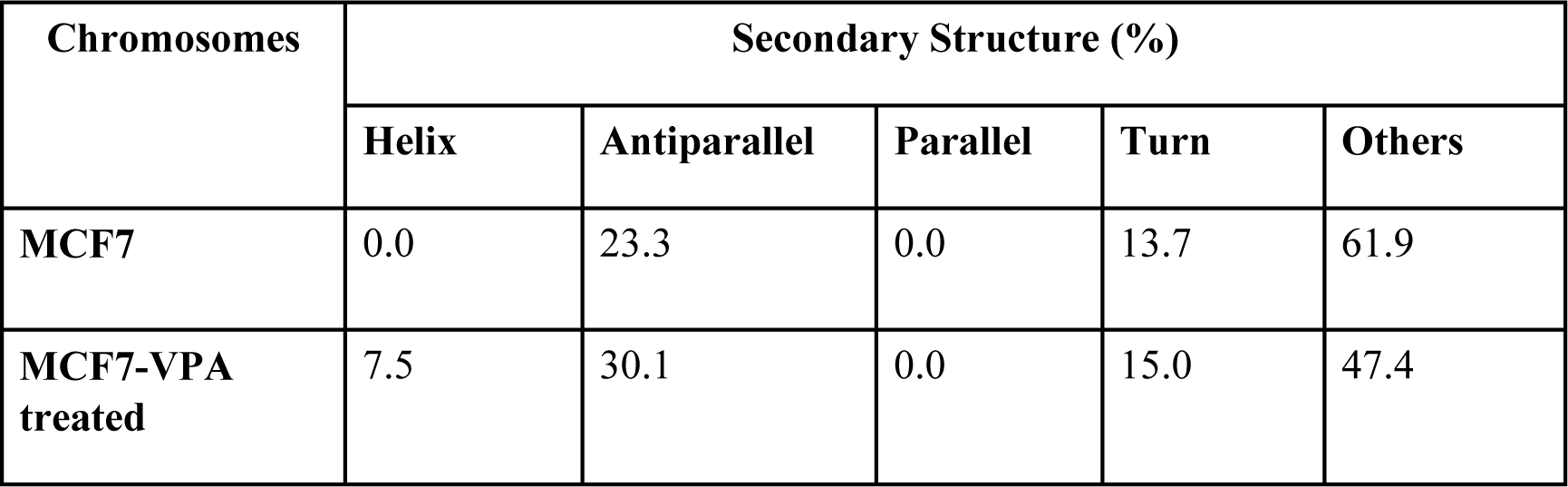
Secondary Structure Calculation from CD Spectra of MCF7 and MCF7-VPA treated chromosomes at room temperature.

| Chromosomes | Secondary Structure (%) |  |  |  |  |
| --- | --- | --- | --- | --- | --- |
|  | Helix | Antiparallel | Parallel | Turn | Others |
| **MCF7** | 0.0 | 23.3 | 0.0 | 13.7 | 61.9 |
| **MCF7-VPA treated** | 7.5 | 30.1 | 0.0 | 15.0 | 47.4 |

**Table S4:** Average roughness, height and Young’s modulus calculation of MCF7 and MCF7-VPA treated chromosomes at different chromosomal regions.

|  | Roughness (nm) | Height (nm) |  |  |  |  | Young's Modulus (MPa) |  |  |  |  |
| --- | --- | --- | --- | --- | --- | --- | --- | --- | --- | --- | --- |
| Chromosomes | Average | Centromere | Telomere | p-arm | Q-arm | Average | Centromere | Telomere | P-arm | Q-arm | Average |
| MCF7 | 18.3 ± 6 | 140 ± 12 | 135 ± 9 | 145 ± 7 | 142 ± 5 | 145 ± 3 | 11.3 ± 3 | 12.8 ± 3 | 12.6 ± 2 | 13.0 ± 2 | 12.5 ± 2 |
| MCF7-VPA treated | 31.9 ± 10 | 125 ± 15 | 121 ± 12 | 128 ± 8 | 120 ± 13 | 134 ± 16 | 1.9 ± 0.8 | 1.9 ± 0.4 | 2.3 ± 0.8 | 2.3 ± 0.7 | 2.0 ± 0.6 |

**Table S5:**
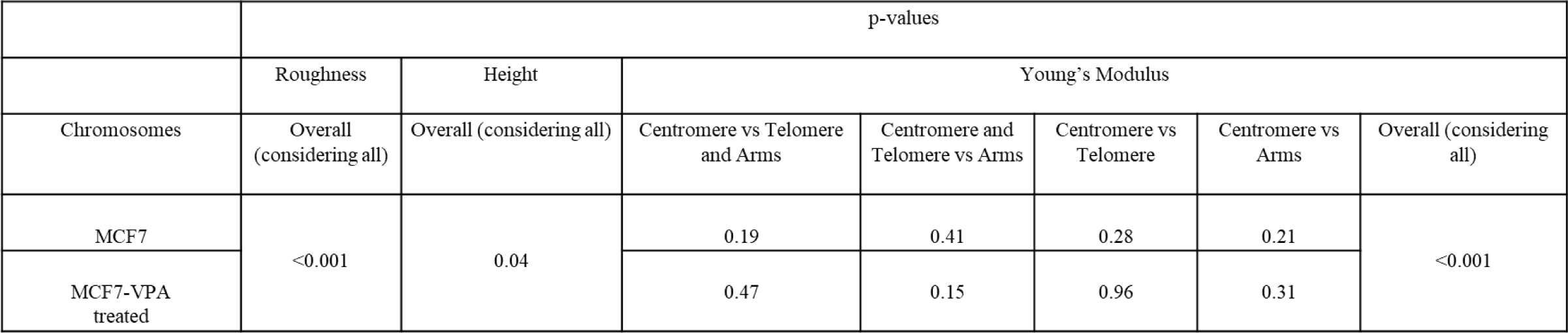
Two-tailed t-test values of Average roughness, height and Young’s modulus of MCF7 and MCF7-VPA treated chromosomes.

**Table S6:** Fowler–Nordheim (F–N) ‘‘B’’ values (V) at different applied force loads for MCF7 and MCF7-VPA treated chromosomes. Two-tailed t-test values of current at applied forces 10, 40 and 80 nN.

| Chromosomes | B-values (V) |  |  | p-values (at 3V) |  |  |
| --- | --- | --- | --- | --- | --- | --- |
|  | 10nN | 40nN | 80nN | 10nN | 40nN | 80nN |
| **MCF7** | 9.85 ± 0.7 | 8.18 ± 0.3 | 15.55 ± 1.6 | <0.01 | <0.01 | <0.01 |
| **MCF7-VPA treated** | 4.40 ± 0.1 | 3.49 ± 0.4 | 6.86 ± 0.3 |  |  |  |

